# Random Forest Regression models for Lactation and Successful Insemination in Holstein Friesian cows

**DOI:** 10.1101/2020.11.17.386318

**Authors:** Lillian Oluoch, László Stachó, László Viharos, Andor Viharos, Edit Mikó

## Abstract

To overcome well-known difficulties in establishing reliable models based on large data sets, the Random Forest Regression (RFR) method is applied to study economical breeding and milk production of dairy cows. As for the features of RFR, there are several positive experiences in various areas of applications supporting that with RFR one can achieve reliable model predictions for industrial production of any product providing a useful base for decisions. In this study, a data set of a period of ten years including about eighty thousand cows was analysed by means of RFR. Ranking of production control parameters is obtained, the most important explanatory variables are found by computing the variances of the target variable on the sets created during the training phases of the RFR. Predictions are made for the milk production and the conception of the calves with high accuracy on given data and simulations are used to investigate prediction accuracy. This paper is primarily concerned with the mathematical aspects of a forthcoming work focused on the agricultural viewpoints. As for future mathematical research plans, the results will be compared with models based on factor analysis and linear regression.

## Introduction

Reproductive management is a key factor in economic dairy production and poor practice can cause considerable economic loss, mainly because of decreased milk yield per cow per lactation and decreased number of calves per year per cow. However, it is also associated with reduced conception rates [1]. Conception rate is determined by heat detection, the choice of the first insemination time after calving, induction of ovulation, and ovulation synchronization program. As we seek for the best conception rate, it is also worth noting the existence of some environmental features and management practices that would directly affect insemination, thus have adverse effects on reproduction performance. The efficiency, accuracy and timing of artificial insemination (AI) remain a major challenge to improving reproductive and economic efficiencies of many dairy farms [6, 14]. Various studies have shown that regression models are of great importance in addressing the issues around the conception rate. Some of them have been used in prediction of the optimal time of insemination [12]. Probability of conception was analysed using a logistic procedure which uses maximum likelihood method to fit linear logistic regression [5]. However, for the logistic regression, the target variables are assumed to be independent and single valued, yet some data are categorical. Due to these deficiencies, other methods like machine learning procedures are sort to be used to address such problems. Various machine learning algorithms including Bayesian networks, decision trees and in particular random forest algorithms have been used for such tasks. Bayesian networks are mainly suited for small and incomplete data [9] with challenges in discretizing continuous variables and implementing recursive feedback loops [13]. Decision trees, as well as RFRs can be used for classification and regression too. RFR algorithm has been widely utilized due to its ability to accommodate complex relationships. RFR calculations can be trivially parallelized, so they can be done on multiple cores of the same CPU. Additionally, the RFR algorithm involves very few statistical assumptions and its hyperparameters can be used to reduce overfitting. The performance of RFR can be explained by the power of ensemble methods to generate high-performance regressors by training a collection of individual regressors. RFR was considered in a study of predicting pregnant versus non pregnant cows at the time of insemination and it proved to be significantly better than other machine learning techniques in [17]. Random forest was also used in attempt to predict conception outcome in dairy cows [17]. On the other hand, mathematical models for the lactation are not new either. Models of lactation curves were early referenced by [3], but due to limitation of the computers and computational difficulties experienced by then, the early models were based on simple logarithmic transformations of exponentials, polynomials, and other linear functions [15]. Another study gave an overview of the parametric models used to fit of lactation curves in dairy cattle by considering linear and non-linear functions [10]. Machine learning approaches have also proved to be vital in the lactation study. Different models based on machine learning in both non-autoregressive and autoregressive cases have been investigated in [16] exhibiting the best performance for both cases with the random forest algorithm. Regression trees have been used in the past to analyse different factors affecting lactation. Researches on effects of the dry period, the lactation parity, the farm, the calving season, the age of the cow, the year of calving and the calving interval have been performed by several authors [4, 11].

Though, previous studies have used other machine learning based models (including RFR) to predict lactation and successful insemination, the proposed study will adopt RFR technique for the same but with different variables. Our purpose is to investigate how the large collection of data gathered in the last ten years used in milk production factories throughout Europe could effectively be analyzed. Therefore, the aim of this study is to apply random forest regression model to predict factors influencing lactation and the success of insemination (SI) as well as the choice of the time of insemination attempts.

## Materials and methods

For this analysis, a large data set was obtained. However, some data were not useful as some information was missing hence were omitted in the study. All data editing and analyses were conducted in the Python, where *pandas* was used for data preparation and scikit-learn, an open source machine learning library. We also used the open source *fastai* library developed at Stanford University (https://github.com/fastai/fastai). In this study we received a dataset from the Agricultural Department of the University of Szeged collected from three major livestock farms concerning 21 different parameters of 82564 Holstein Friesian cows. In this case we considered the variables subdivided into three groups as follows:

V1. *Genealogy:* (i) Settlement (ST in later abbreviation); (ii) Cow ID (ID); (iii) Father (FT); (iv) Parity (Calving Number, CN); (v) Calving Date (CDT); (vi) Sex of 1st Calf (SX1); (vii) Sex of 2nd Calf (SX2).

V2. *Insemination and calving:* (i) First Date of Insemination after Calving (FDFB); (ii) Time between calving and first insemination (DFI); (iii) Date of successful insemination after calving (PSIS); (iv) Time between calving and successful insemination (SFAC); (v) Number of unsuccessful inseminations after calving (NUIC); (vi) How many inseminations were unsuccessful in the previous calving (IPAC); (vii) Days open (Number of days to successful insemination in previous lactation, PCIS); (viii) Age in months of conception at heifer (UMP); (ix) Age in months at first calving (MFCA).

V3. *Lactation:* (i) Milk yield in previous lactation (AMPL); (ii) 305-day Milk Yield in previous lactation (AMPLD); (iii) Number of days milking previous lactation (DPLM); (iv) Dry days in previous lactation (PLD); (v) Calving interval (DC).

Notice that the data structure was not intended to be the subject of a deep mathematical analysis. Furthermore, it is easy to see that there are parameters with obviously high correlation, on the other hand it would have been more useful to have an access of genuine time series in details instead of accumulated parameters like the calving interval. It is also remarkable that the current data do not involve finer details of production volumes of lactation. Studies concerning lactation and BLUP index [8] will be done in a separate work to be submitted in an agricultural journal. Here we concentrate on the description of the use of RFR techniques and we restrict our attention to the following problems:

P1. To estimate the 305-day milk yield in previous lactation on the basis of the variables except for 3(ii).

P2. To estimate the number of days open on the basis of the variables except for 2(iv).

P3. To provide a weighted ranking of the impact of the variables in the answer of the above two questions.

For the targets of the problems P1 and P2, we built respective random forests consisting of 1000 decision trees each. The trees are constructed using the familiar CART algorithm with respect to the hyperparameters (https://doi.org/10.1023/A:1010933404324). Each tree is trained on a bootstrap set. The number of samples drawn from the processed training set to form these bootstrap sets equals the number of samples in the processed training set. The number of samples drawn is a hyperparameter. The maximum depth of the trees is also hyperparameter. We did not limit the depth. Figure 1. and Figure 2. depict sample decision trees of depth 3. The depth of the trees used in the models is much bigger. For each tree, there is one node at the start, the root node, that contains all the samples. For each node that has at least 2 samples, a split is performed. For each node to be split, the CART algorithm examines the possible splits of all the features for that specific node and the best alternative is considered, according to the used splitting criterion. Subsequently, the interval of the selected feature will be split by the selected value of the feature, resulting in two new nodes. Consequently, the two new nodes will have fewer samples. Then, if one or both of these new nodes have at least 2 samples, the splitting process continues. When there is only 1 sample left in a node, the CART algorithm will not perform the splitting. The minimum number of samples required for splitting is also a hyperparameter and can be changed. We used the default value, 2. The trees are grown as long as no stopping rule stops the growing process. Pure nodes, where the target variable is identical in all samples, are not split.Nonetheless, we did not use pruning. This way, we got “fully grown and unpruned trees”(https://scikit-learn.org/stable/modules/generated/sklearn.ensemble.RandomForestRegressor.html. and https://doi.org/10.1023/A:1010933404324).

**Fig 1.**
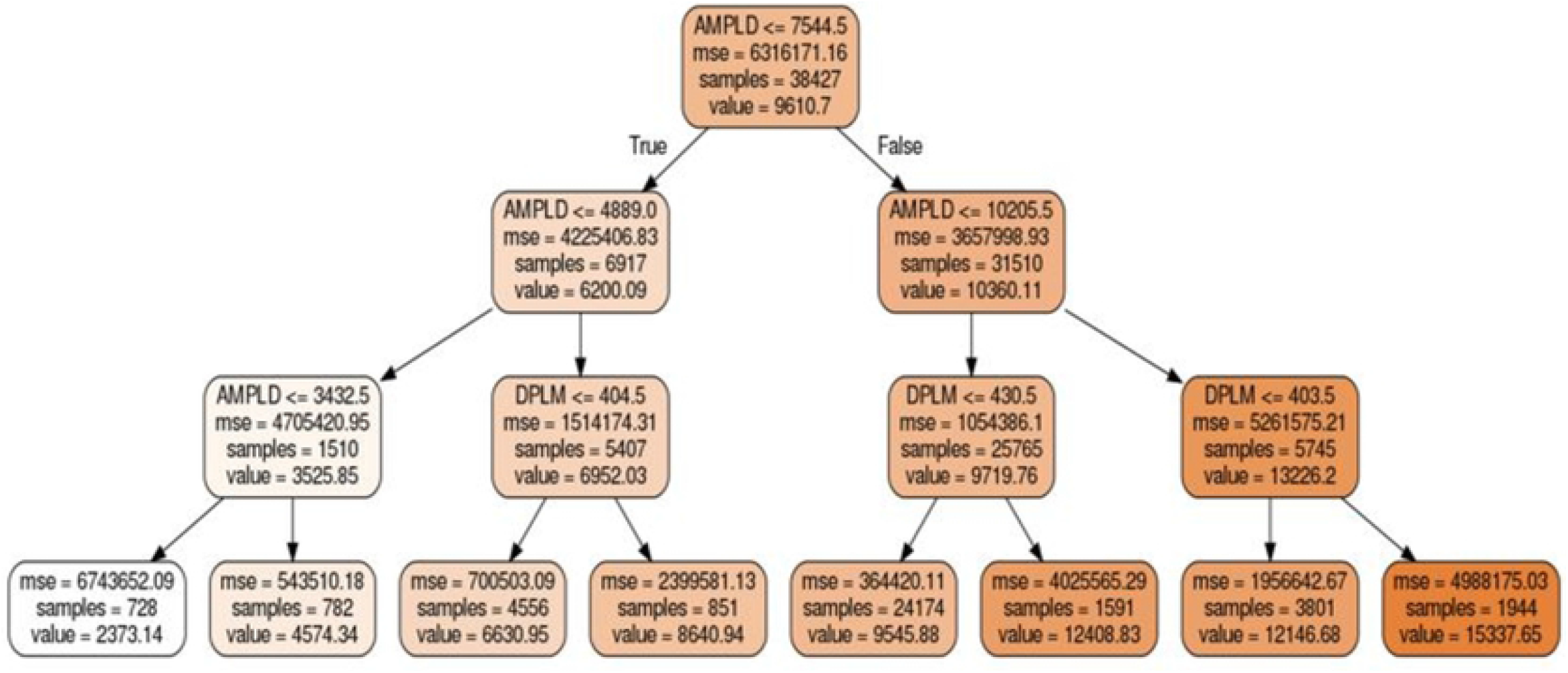
Lactation random tree. Sample of unpruned random tree for lactation taken from the data of 82564 Holstein Friesian cows

**Fig 2.**
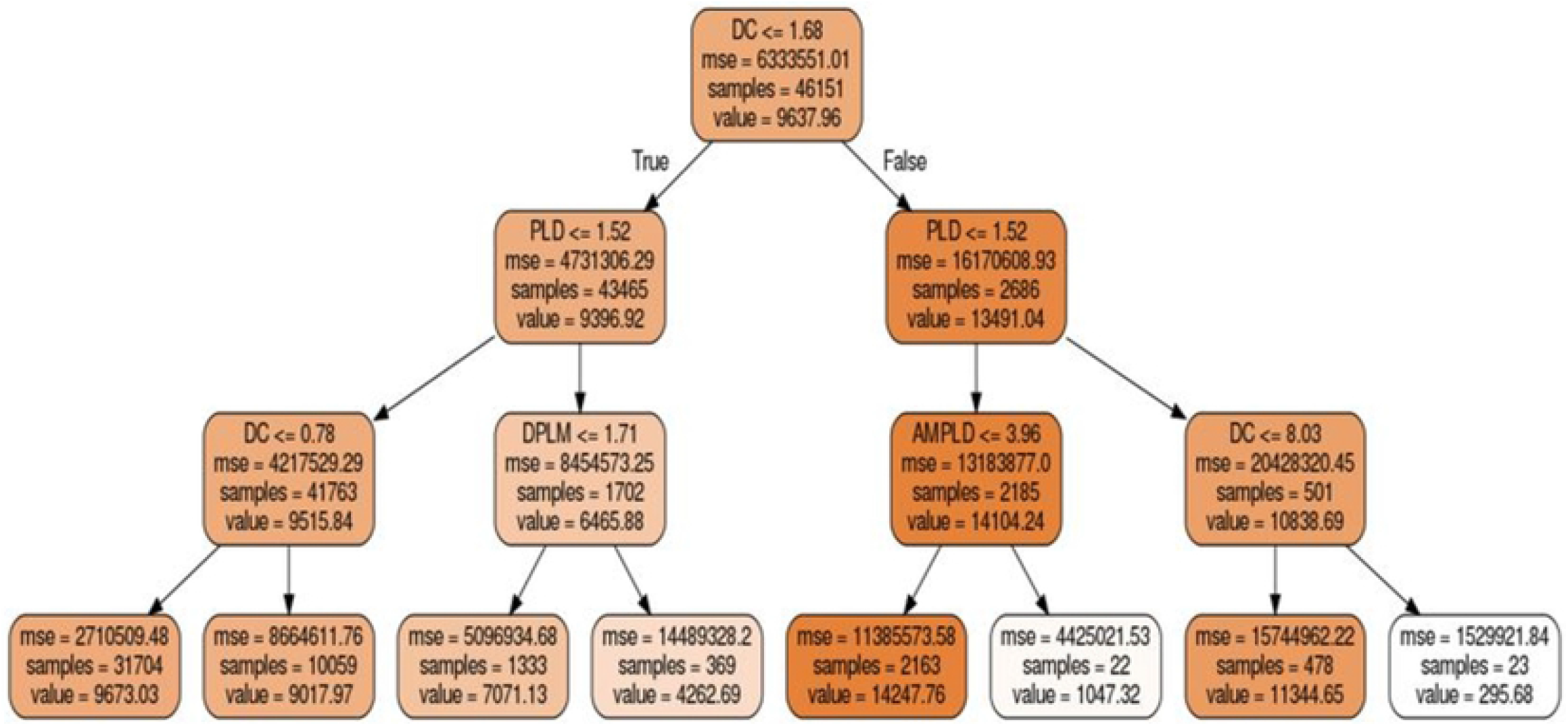
Successful insemination random tree. Sample of unpruned random tree for Successful Insemination taken from the data of 82564 Holstein Friesian cows.

In both cases, the prediction value by means of the random forest is the mean of the prediction values of its individual decision trees.

Given any tree *T* and one of it nodes *n*, our procedure fixes a feature *f_T,n_* along with a value *v_T,n_* in the range of *f_T,n_*. Furthermore the algorithm fixes a prediction value *p_T,l_* to every leaf of *T*. Given a ‘‘virtual cow” *C* with *f_T,n_*-values *f_T,n_*(*C*) (for each node *n* of *T*), our RFR estimate for answering P1. P2 according to tree *T* is the value *r_T_*(*C*) = *p_T,l_*(*c*) where the leaf *l*(*C*) is determined as follows: We start at the root of the tree, and if a node *n* is reached, the decision for continuing to left or right to a next node is done accordingly, if we have *f_T,n_*(*C*) < *v_T,n_* or *f_T,n_*(*C*) ≥ *v_T,n_* (cf. Fig 1 and Fig 2). Our steps end when reaching a leaf which we set to *l*(*C*).

The prediction value by means of a random forest is the mean of the prediction values of its individual decision trees. As for standard theoretical background we refer to and the monograph [7] and Breiman [2]. For implementation we used the python machine learning package scikit-learn (https://scikit-learn.org/stable/). As we know, scikit-learn’s RandomForestRegressor object is trained using the CART algorithm.

After the training is done, we need to check its score on the test set. If the *R*^2^ score of the forest’s predictions on the test set is not good, then we need to fine-tune the hyperparameters of the RFR. When using hyperparameter tuning, we make a grid of the hyperparameters of the RFR (maximum depth of the trees, minimum samples for splitting, change pruning, etc.). Additionally, we need to split the training set into a real training set and a validation set. We will use the latter to find the best set of hyperparemeters. Then we need to train a RFR for each value in the grid, and check their predictive performance using cross-validation. Consequently, the best set of hyperparameters is used for our final model choice. We then train the RFR on the combined training and validation sets, and check its final predictive performance on the test set, that we did not use at until this final step.

If the model performs well, we can decide to not deal with hyperparameter tuning, because the model is already good enough.

The original data set consists of 82,563 records, but it contains missing data and invalid data. For this reason, not all records were kept. As for the Lactation model, 45,461 records were kept, as for the Succesful Insemination model, 82,378 records were kept after the preparation of the data set. In our study, we used 10% of the prepared data set as the test set, and the remainining 90% of the data set as the training set. Actually we ended up having a good model for the first time with great *R*^2^ scores on the test set. Hence we did not need hyperparameter tuning and a separate validation set. Of course, hyperparameter tuning could still be done. It could still improve the model by a small margin, but we did not think we needed to do this.

It is remarkable that the RandomForestRegressor object of scikit-learn automatically calculates the feature importance values of the variables. The sum of these feature importances is 1. As for the case of artificial insemination, there are two features that stand out with feature importance values of 0.544 and 0.255. For lactation, there are two features with importances 0.88 and 0.084. Hence we can investigate the effect of the two most important explanatory variables by keeping the remaining variables at their median values.

## Results and Discussion

Our related program files and the detailed outputs are deposited on the departmental webpage (http://www.mgk.u-szeged.hu/karunkrol/kutatas/cow-article). Although random forest models can handle correlation among the data well, we investigated the dataset in this aspect too.

Figures 3(a) and 3(b) presents the outcome of correlation coefficient for lactation analysis and successful insemination analysis respectively. It was observed that several variable showed relatively high correlations for the lactation analysis, AMPL and AMPLD (0.88), DPLM and PLESI (0.86), PLESI and DC (0.84) DPLM and DC (0.73). However for SI analysis IPAC and SFAC (0.73) was the only highly correlated case.

**Fig 3.**
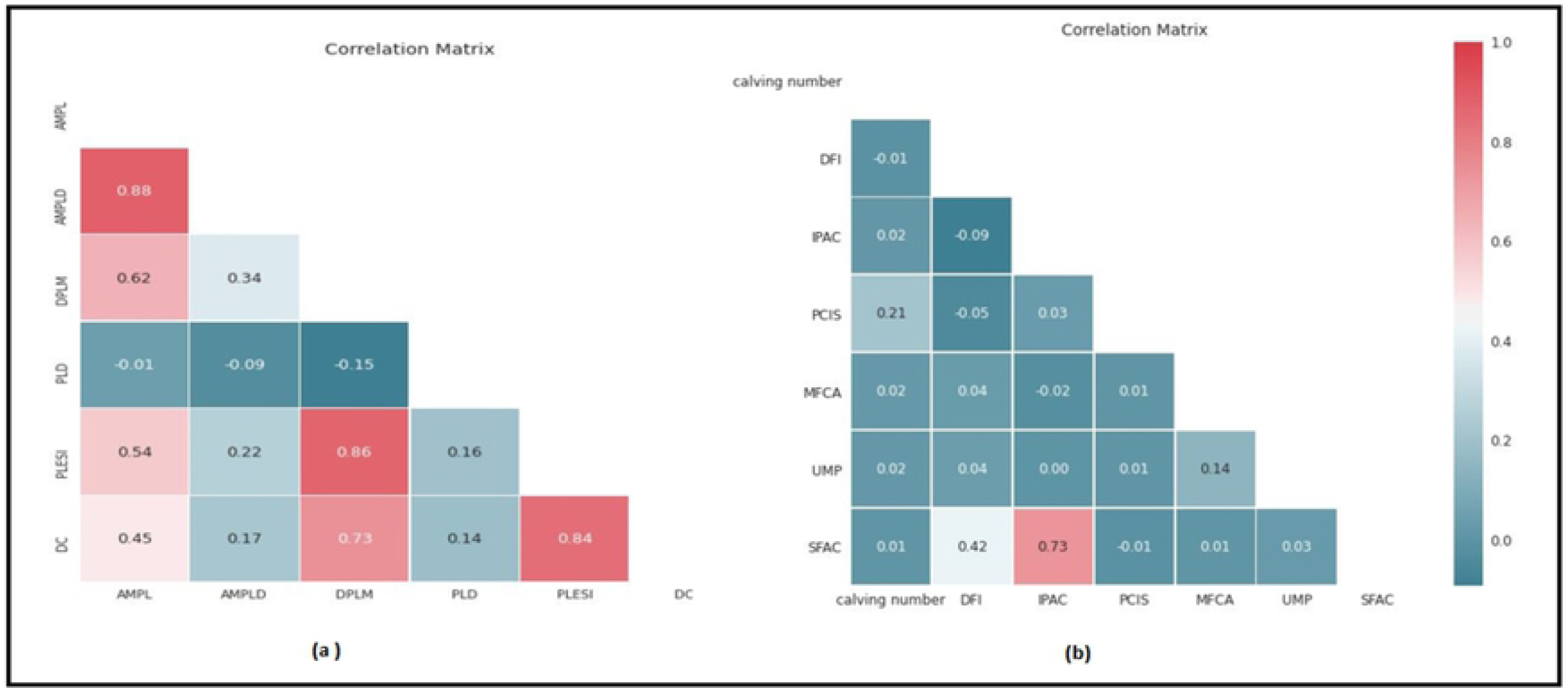
Correlation matrix for Lactation and Successful Insemination. Correlation values between various variables for(a) Lactation and (b) Successful Insemination

Separation of data into the features and targets was achieved by first splitting data into training and testing sets. It is expected that there would be some relationship between all the features and the target value, and so the model learns this relationship during training. With regards to quantifying the predictive information provided by the variables in the entire random forest, the feature importance of the variables plays an important role. It was evident that AMPLD plays a vital role in the Lactation model with a feature importance of 88.0%, followed by DPLM (8.4%). On the other hand the most important predictor variable for SI model was IPAC(54.4%) and DFI (25.5%). Figure 4(a) and 4(b) indicates the feature importance of the explanatory variables in relation to the target variable for lactation and successful inseminations respectively.

**Fig 4.**
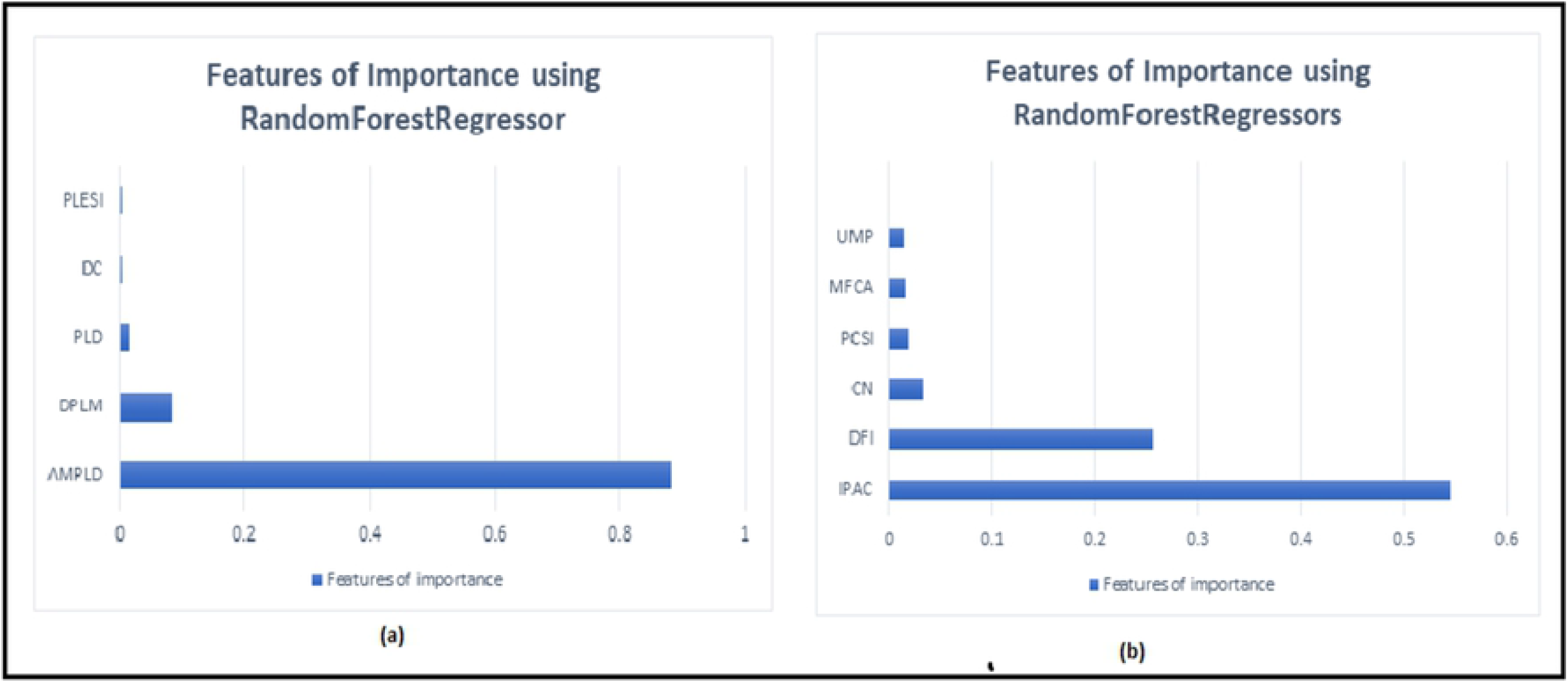
Features of importance for Lactation and Successful Insemination. Features importance of variables for (a) Lactation and (b) Successful Insemination in percentage.

Creating and training the model simply involves instantiation of the program object RandomForestRegressor and fitting it on the training data. This results to a large number of expansive trees that forms the forest (1000 trees in this case), which are essential in making and evaluating the reasonable predictions. The goodness of fit can be evaluated by means of the *R*^2^ values shown in Table 1.

**Table 1.** *R*^2^ for training and test set for Lactation and SI

|  | Training set | Test set |
| --- | --- | --- |
| Lactation | 0.998 | 0.987 |
| SI | 0.993 | 0.948 |

Since the model performed well on the test set, we were not forced to fine-tune its hyperparameters and use a validation set. Fine-tuning of the hyperparameters and checking the model performance on a separate validation set is likely to further improve the prediction accuracy of our model. This is a possibility to develop the model.

Based on the results from features of importance of the random regressor for both the Lactation model and SI model the results of the model are summarized in Tables 2 and Table 3.

**Table 2.** Features of importances by the random regressor for the Lactation model

| Predictors | AMPLD | DPLM | PLD | DC | PLESI |
| --- | --- | --- | --- | --- | --- |
| Model Parameters | 0.880 | 0.084 | 0.015 | 0.004 | 0.003 |

**Table 3.**
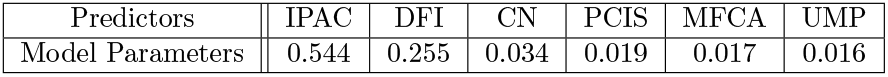
Features of importances by the random regressor for SI

For both target variables, the two most important explanatory variables have significantly higher feature importance values than the rest of the variables. We tried to visualise the way these affect the targets. We made a data set for each target variable by varying both of the two most important explanatory variables while keeping the rest of the variables at their median values. Then we used the trained random forest models to predict the target variables. The results are illustrated in function graph diagrams shown in the concluding Figures 5(a) and 5(b). This way, we can get an idea about the way these two most important variables affect the target variables. The actual effects cannot be visualised in 3D graphs because of the interactions with all the other variables. These graphs can be analysed for designing production strategies without specific mathematical preparation by agricultural experts.

**Fig 5.**
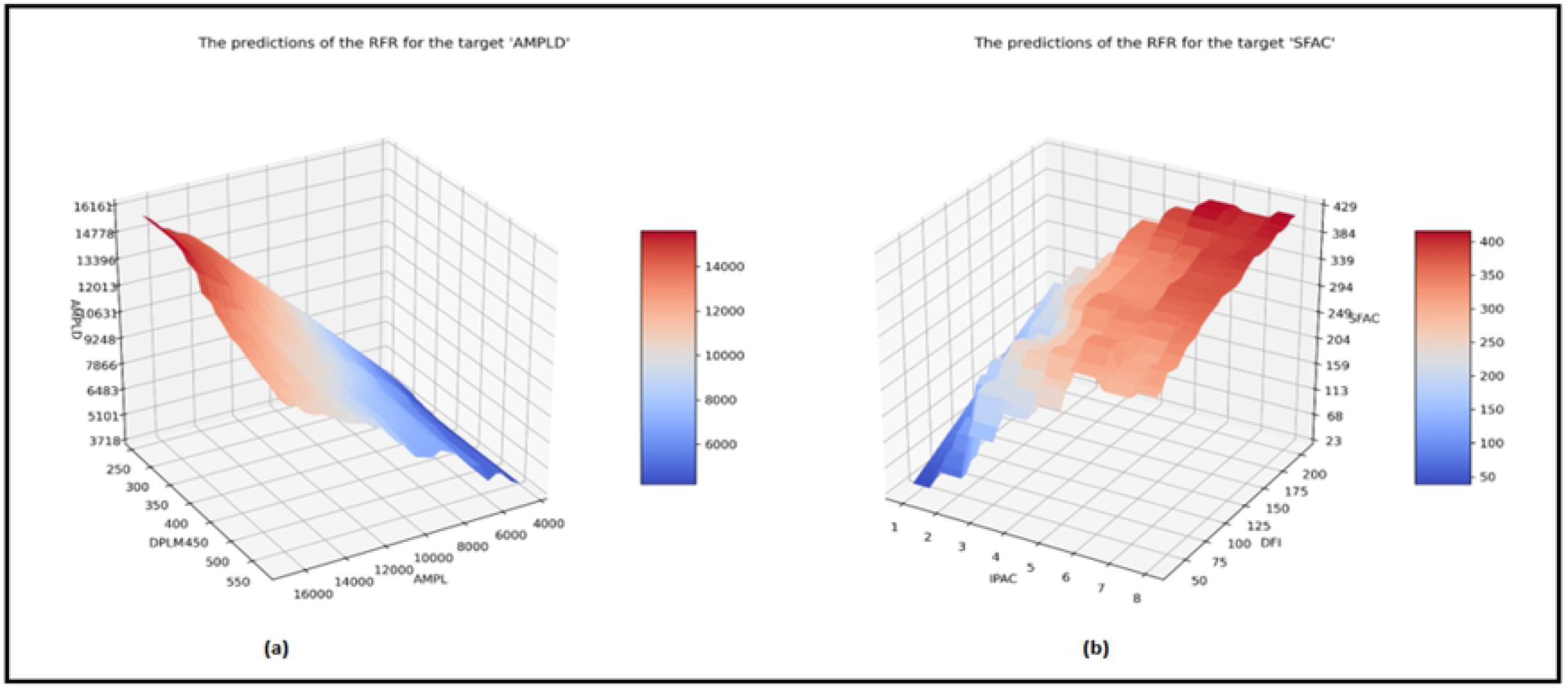
RandomForestRegressor predictions for Lactation and Successful Insemination. Diagram for the Predictions of RFR for (a) Lactation and (b) Successful Insemination.

## Conclusion

We established an alternative approach to other machine learning based models concerning Questions P1,P2,P3 by means of random forest regression. The transformed dataset were split into a test size of 10%, the remaining 90% was used to train the forest. The results indicated that when the target was *SI*, the prediction of the RFR on the test set and the actual targets had an *R*^2^ ≈ 0.948. The important features were the IPAC, DFI, CN, PCIS, MFCA and UMP with importance scores in Table 3. When the target was *Lactation*, the prediction of the RFR on the test set and the actual targets had an *R*^2^ ≈ 0.987. The important features of were AMPLD, DPLM, PLD, DC and PLESI with importance scores in Table 2. Various alternative regression methods were used for analysis of this data set but all these attempts failed due to the complexity of data and the large sample size. It seems that the RFR is good for practical applications (as for problems P1, P2 and P3.)

## Acknowledgments

This research was supported by the Ministry of Human Capacities, Hungary grant TUDFO/47138-1/2019-ITM.The research due to the fourth author was partially carried out at BME and has been supported by the NRDI Fund (TKP2020 IES, Grant No. BME-IE-MISC) based on the charter of bolster issued by the NRDI Office under the auspices of the Ministry for Innovation and Technology.

## Authors Contribution

L.S, L.V and E.M conceived of the presented idea. L.O and A.V processed the data, performed the analysis and designed the figures. E.M contributed the analysed data. L.O took the lead in writing the manuscript. All authors provided critical feedback and helped shape the research, analysis and manuscript. L.S supervised the project.

